# Sex-specific effects of fatiguing exercise on skeletal muscle passive mechanics are preserved in aging

**DOI:** 10.64898/2026.05.22.727297

**Authors:** Grace E. Privett, Julissa Ortiz-Delatorre, Austin W. Ricci, Karen Wiedenfeld Needham, Damien M. Callahan

## Abstract

Skeletal muscle function is central to the preservation of functional mobility. Given global shifts to an increasingly aged population, it is paramount that researchers and clinicians better understand the effectors of age-related functional decline. Muscle fatiguability acutely modifies skeletal muscle mechanics in ways that may affect joint stability. We have previously reported sex-specific reductions in cellular passive stress and modulus with fatigue in young males, but not females. Here, we assess whether older adults, who are more susceptible to fatigue during dynamic contractions, exhibit changes to cellular passive mechanics following fatiguing exercise. Muscle tissue biopsies were collected from 11 young and 11 older adults to measure passive stress and Young’s Modulus at the single fiber and bundle level. Biopsy samples were acquired from rested muscle and immediately following intermittent maximal contractions to task failure. Fatigue was associated with persistent reduction in elastic modulus that was specific to male participants, regardless of age. In muscle fiber bundles, containing both myofibrillar proteins and the extracellular matrix, fatigue-induced changes in modulus were largely negated, with the only significant change observed in young females, who demonstrated enhanced modulus with fatigue. Taken together our findings suggest a preservation of sex-based differences in the acute response to fatigue across the adult lifespan when measured at the myofilament level. However, further research is needed to understand how and whether these findings translate to the whole tissue level.

**New and noteworthy:** Acute modifications to muscle tissue mechanics are poorly understood but may have important impacts on functional outcomes in at-risk populations. Our findings suggest myocellular mechanics respond to acute fatigue stress in a sex specific manner that persists across the lifespan.

## Introduction

The global population of older adults is rapidly expanding (1–3), exacerbating the burden of age-related morbidity on healthcare systems. Aging is associated with the progressive loss of skeletal muscle strength and mass, presenting a clear threat to mobility and independence of living in older adults. Skeletal muscle stiffness, the tendency to resist deformation in response to an applied force, is increased in aging due to increases in extracellular matrix (ECM) stiffness and intramuscular accumulation of connective and adipose tissue (4). Increased muscle stiffness, along with antagonist coactivation, limits joint mobility and angular velocity (5) thereby hindering activities of daily living.

Advanced age is accompanied by increased fatiguability during repeated dynamic tasks (6–8). Because fatigue is linked to increased falls risk in this population (9), it is important to understand age-related differences in tissue stiffness responses to the physiological stress of repeated contractions. Previous studies have reported reduced skeletal muscle passive stiffness following fatiguing exercise in whole muscle (10, 11) and single fibers (12, 13), which may contribute to fatigue-induced falls via diminished joint stability (14) or translation of myocellular force to the tendon. Within muscle fibers, passive mechanical properties derive from the viscoelastic protein titin (15, 16); therefore, fatigue-induced reductions to cellular measures of stiffness (i.e. Young’s Modulus, stress) likely involve titin-related mechanisms including titin post-translational modifications resulting from fatiguing exercise or age-related neuromuscular changes. However, whether fatiguing exercise impacts skeletal muscle measures of stiffness in older adults has not been previously studied. Therefore, we sought to extend our previous work (12, 13) by exploring the relationship between acute fatigue and skeletal muscle passive mechanics in older adults. We extended our investigations of myocellular mechanics to the tissue level by assessing elastic modulus in bundles of muscle tissue, containing both single fibers and associated ECM, to consider the influence of age-related increase in ECM stiffness on the present measures (17). While the primary effects of age on passive muscle tissue modulus have been explored, the potential impacts of acute, exercise-induced fatigue have not yet been studied. Together, our single fiber and bundle level assessments of passive tissue mechanics provide an intriguing assessment of the chronic and acute changes that occur to skeletal muscle mechanical properties at the cellular and tissue levels.

Age-related attenuation of skeletal muscle excitation-contraction coupling and beta-adrenergic signaling, coupled with reported increases in muscle fiber stiffness with age (16, 18), support the hypothesis that measures of cellular passive stress and modulus would be higher in non-fatigued fibers from older versus younger adults, and that fatigue-induced compliance would be less in fibers from older versus younger adults. Similarly, age-related increases in skeletal muscle tissue-level (4, 5) stiffness support the hypothesis that passive modulus would be higher, and fatigue would induce less compliance in bundles from older versus younger adults. Such findings would reflect an impaired capacity to adjust cellular and tissue mechanics to length perturbations following fatiguing exercise in older adults. If present, this age-related alteration in tissue mechanics may contribute to increased falls risk at fatigue in older adults.

## Methods

### Population

This protocol was approved by the Institutional Review Board at the University of Oregon. 11 older (aged 65-80) and 11 younger (18-35 years) males and females from the University of Oregon and surrounding community consented to participate in this study. ActivePal accelerometers (Glasgow, Scotland) were affixed to the outer-mid thigh with an adhesive in order to characterize physical activity levels in the present cohort (19). Physical activity was monitored for an average of 8 ± 2 days, which exceeds the recommended 3 days (20). To limit the potential for menstrual cycle-dependent variation in circulating estradiol and associated potential impacts on skeletal muscle mechanical properties, all female volunteers either reported use of hormonal contraceptive or were tested in the pre-follicular phase of the menstrual cycle, (within 5 days of menses onset). Participants reported no orthopedic limitations (severe osteoarthritis, joint replacement, or other orthopedic surgery in the previous six months), endocrine disease (hypo/hyper thyroidism, Addison’s Disease or Cushing’s syndrome), uncontrolled hypertension (>140/90 mmHg), neuromuscular disorder, significant heart, liver, kidney or respiratory disease, or diabetes. Participants were non-tobacco-smokers and had no current alcohol disorder. Finally, participants taking medications known to affect muscle stiffness or beta-adrenergic signaling of neuromuscular activation (including but not limited to beta blockers, calcium channel blockers, and muscle relaxers) or anabolic steroids were not included.

### Study Design

Participants visited the lab on 2 occasions separated by at least 1 week. During the first visit, non-invasive measures of voluntary strength, power, and fatigue of their dominant knee extensors (KE) were collected using a Biodex System 3 dynamometer (Biodex Medical Systems, Shirley, NY) and participants were familiarized with the exercise protocol. During the second visit, volunteers performed maximal voluntary isometric contractions of the KE followed by fatiguing exercise to task failure. Fatiguing exercise was followed by bilateral, percutaneous needle muscle biopsies: one on the exercised limb immediately following exercise (“fatigued”) and the second on the contralateral, non-exercised limb (“non-fatigued”).

### Fatigue Protocol

Participants exercised the dominant limb on a Biodex System 3 dynamometer (Biodex Medical Systems, Shirley, NY). Participants were seated on the dynamometer with hips and knee flexed at 90° (180° = full extension). Prior to fatiguing exercise, participants completed three maximum voluntary isometric contractions (MVIC) of the knee extensors while analog voltage data for torque were sampled at 500 Hz. These initial 3 MVICs were used to quantify absolute and relative peak torque, power, and velocity of the dominant knee extensors. Analog data were converted to digital using an analog-to-digital converter (Cambridge Electronic Design, UK). Real-time visual feedback was provided to the participants to encourage maximal effort during the MVIC. The average torque value of the three MVICs was used to set the applied load to 30% MVIC maximal torque for the bout of fatiguing exercise. Following initial MVICs, participants performed repeated, voluntary knee extensions at this isotonic load until task failure. Task failure was identified as the inability to perform knee extension through at least 50% range of motion. Fatigue was quantified as the Fatigue 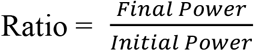, where “initial power” represents the average peak power of the first five knee extensions performed during fatiguing exercise, and “final power” represents the average peak power from the last five knee extensions. Time to fatigue (task failure) was recorded for all participants.

### Muscle Biopsy Procedure

Percutaneous needle biopsy of the VL was performed within 10 ± 4 minutes following task failure, under sterile conditions as previously detailed (21). First, the biopsy site was sterilized and local anesthetic (1 or 2% lidocaine HCL [Hospira Worldwide, Lake Forest, IL, USA]) was administered via injection. Next, a small (~5 mm) incision was made in the skin and muscle fascia, allowing for the passage of a Bergstrom biopsy needle (5 mm diameter) to the belly of the VL muscle to acquire sample at a depth of ~2-3 cm. Following acquisition of biopsy sample, the collected muscle was retrieved from the needle using forceps. Time elapsed before acquisition of a biopsy sample was collected for 7/8 older adults and 6/9 younger adults. Average time elapsed = 11 ± 3 minutes in older adults and 9 ± 5 minutes in younger adults.

### Tissue Processing and Dissection

Sample collected from the biopsy was placed in muscle dissecting solution (MDS, 120.782 mM sodium methanesulfonate (NaMS), 5.00 mM EGTA, 0.118 mM CaCl_2_, 1.00 mM MgCl_2_, 5.00 mM ATP-Na_2_H_2_, 0.25 mM KH_2_PO_4_, 20.00 mM BES, 1.789 mM KOH, 1 mM Dithiothreitol (DTT)), parsed into bundles of ~50 fibers, and tied to glass rods before advancement through solutions of increasing glycerol content and storage in 50% glycerol solution (5.00 mM EGTA, 2.50 mM MgCl_2_, 2.50 mM ATP-Na_2_H_2_, 10 mM imidazole, 170.00 mM potassium propionate, 1.00 mM sodium azide, 50% glycerol by volume) at −20°C. Sample allocated for mechanics analyses were used within 4 weeks following biopsy. For permeabilized fiber experiments, one glycerol-stored bundle was chemically skinned (“secondary skinning”, MDS + 1% Triton X-100) prior to fiber dissection. Dissected fibers were then chemically skinned, transferred to plain MDS, and kept on ice until experimentation. For bundle experiments, glycerol-stored bundles were not subjected to secondary skinning. One glycerol-stored bundle was placed directly into plain MDS, then groups of 10-20 fibers (~350 µm diameter) with intact ECM were dissected and stored on ice until experimentation.

### Single fiber and bundle morphology and contractile measures

Prepared fibers and bundles were mounted in relaxing solution (67.286 mM NaMS, 5.00 mM EGTA, 0.118 mM CaCl_2_, 6.867 mM MgCl_2_, 0.25 mM KH_2_PO_4_, 20.00 mM BES, 0.262 mM KOH, 1.00 mM DTT, 5.392 mM Mg-ATP, 15.00 mM creatine phosphate (CP), 300 U/mL creatine phosphokinase (CPK)) between a force transducer and a length motor (Aurora Scientific, Inc., Aurora, ON, Canada). Single fibers were mounted using the Moss clamp technique (22). Bundles were measured by direct suturing of the bundle to the rods attached to the force transducer and length motor. Fiber and bundle dimensions were measured as follows: d_top_ = average of three diameter measures along the fiber using the top-down view; d_side_ = average of three diameter measures along the fiber using the side view; fiber length = distance between the two trough edges. Passive tension was measured by zeroing the force transducer while the fiber or bundle was slacked, then stretching the sample to SL = 2.65 µm (single fibers) or SL = 2.40 µm (bundles). In single fibers, active tension was measured at SL 2.65 µm by moving the fiber to pre-activating solution (81.181 mM NaMS, 5.00 mM EGTA, 0.012 mM CaCl_2_, 6.724 mM MgCl_2_, 5.00 mM KH_2_PO_4_, 20.00 mM BES, 1.00 mM DTT, 5.397 mM Mg-ATP, 15.00 mM CP, 300 U/mL CPK) followed by activating solution (57.549 mM NaMS, 5.00 mM EGTA, 5.021 mM CaCl_2_, 6.711 mM MgCl_2_, 5.00 mM KH_2_PO_4_, 20.00 mM BES, 9.674 mM KOH, 1.00 mM DTT, 5.437 mM Mg-ATP, 15.00 mM CP, 300 U/mL CPK). Once a steady state tension was recorded, the sample was returned to relaxing solution. All fibers were activated (pCa 4.5) prior to passive stretching to measure active tension and confirm fiber viability. Bundles were not activated prior to passive stretch measures.

### Passive stretch protocol

For both fibers and bundles, passive stretch measures were performed in relaxing solution (pCa 8.0) using a passive stretch protocol adapted from previous work (16), as described previously (12). Initial sarcomere length (SL) was set to 2.4 µm, followed by 7 incremental stretches to reach a final length of 156% of initial length (SL ~3.8 µm). Each stretch lengthened the sample 8% of initial length in 2 seconds and held this position for 2 minutes of stress-relaxation. SL was measured throughout the protocol using an inverted microscope located beneath the single fiber rig. Following completion of the passive stretch protocol, each fiber was collected and placed in gel loading buffer (2% SDS, 62.5 mM Tris, 10% glycerol, 0.001% bromophenol blue, 5% β-mercaptoethanol, pH 6.8), centrifuged and heated at 65ºC for 2 minutes, then stored at −80°C until later assessment of myosin heavy chain (MHC) isoform.

### MHC isoform identification

Sodium dodecyl sulfate poly acrylamide gel electrophoresis (SDS-PAGE) was used to determine the MHC isoform of single muscle fibers. Sample from each fiber was loaded into its own well of a 4% stacking / 7% resolving polyacrylamide gel. The gel was run at 70 V for 3.5 hours followed by 200 V for 20 hours at 4°C (23). Gels were stained with silver and the resulting MHC isoform (I, IIA, and/or IIX) expression was determined by comparison to a standard made from a multi-fiber homogenate.

### Outcome Measures

At the whole-muscle level, relative peak torque and power were calculated by dividing absolute values for torque (N) and power (W) by the participant body weight (kg). For single fibers, maximally activated tension was quantified as the measure of steady-state active force divided by fiber CSA. For fibers and bundles, passive stress at each SL was calculated as 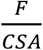, where “F” indicates the measured force value at the end of stress relaxation (Figure 1) and “CSA” indicates fiber or bundle cross sectional area assuming elliptical shape. 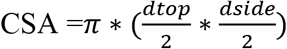 where “d_top_” is the average of three top diameter measures, and “d_side_” is the average of three side diameter measures. Strain was calculated as 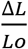, where “L_0_” indicates initial length. Passive stiffness was quantified as Young’s Modulus to account for potential differences in fiber size across samples. For single fibers, passive Young’s Modulus was calculated as the slope of the stress-strain relationship at shorter fiber lengths (strain = 1.00-1.24 %Lo) and at longer fiber lengths (strain = 1.32-1.56 %Lo) to consider the length dependence of cellular passive modulus measures (18). Given the linearity of the stress-strain curve of the bundles, one value for Passive Young’s Modulus was calculated as the slope of the entire stress-strain curve (rather than the “short length” and “long length” values used for single fibers). To consider the potential for variability in the size and number of fibers included in dissected bundles, the ratio of number of fibers to bundle CSA was calculated as the fiber to bundle ratio for each sample.

**Figure 1.**
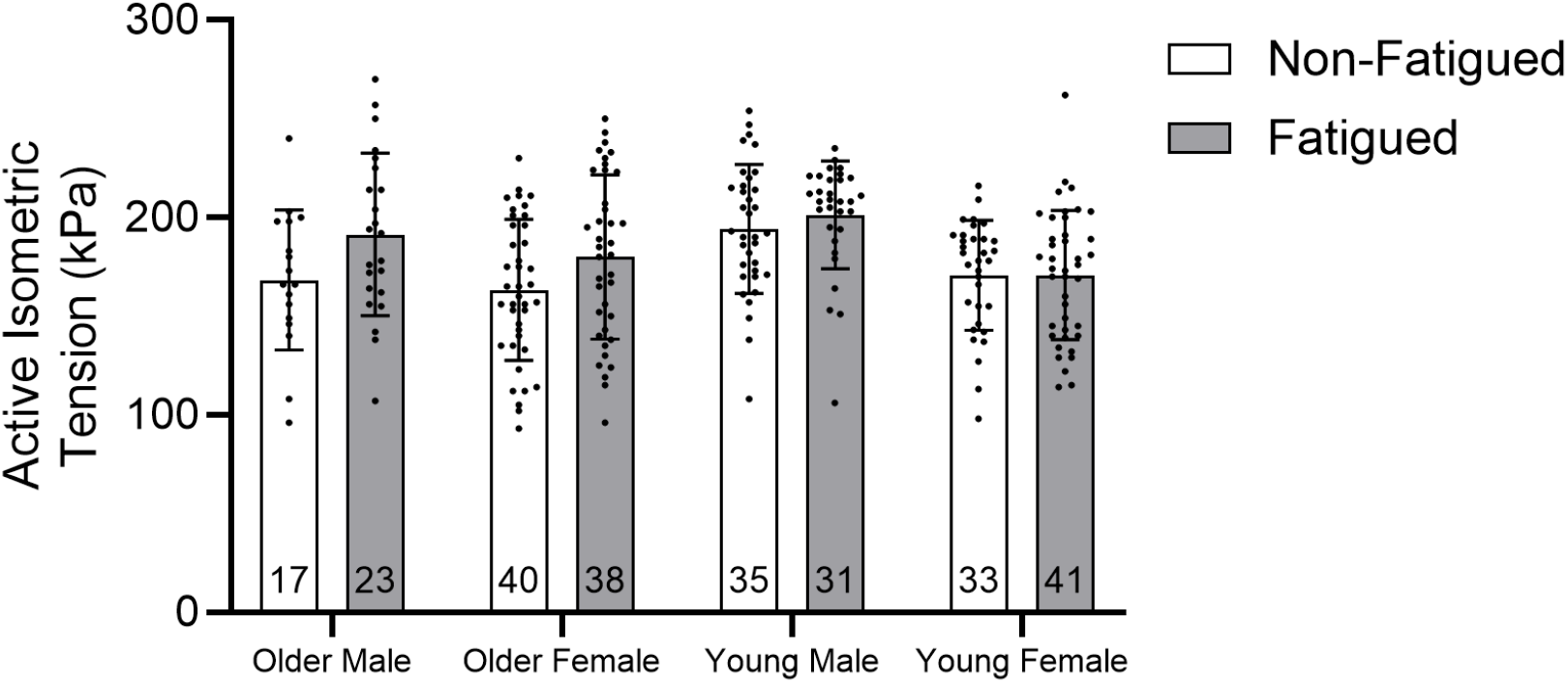
In this sample of MHC IIA and IIA/X fibers, active isometric tension was not affected by fatigue (0.080), biological sex (p=0.410), or age (p=0.569). Data are shown as mean ± SD.

### Statistical Analyses

Statistical testing was conducted using SPSS software package (SPSS, IBM Corp., Armonk, NY, USA), unless otherwise specified. Anthropometric measures and activity data were compared between older and younger males and females using a two-way ANOVA. To evaluate differences in single fiber passive stress and passive modulus (short and long lengths), separate linear mixed models were run with fatigue, age, biological sex, and interaction terms as fixed effects and participant ID as a random effect to account for fiber variation within individuals, as described previously (24). Subsequent analyses to investigate interactions between age and fatigue or biological sex and fatigue used separate linear mixed models in older and younger males and females, each including fatigue as a fixed effect and participant ID as a random effect. To determine whether maximally activated tension, single fiber CSA, or fiber length differed by age, biological sex or fatigue, a linear mixed effects model was run with age, sex, fatigue, and interaction terms as main effects and participant ID as a random effect. To test for significant differences in the passive Young’s Modulus of bundle samples, a linear mixed model was generated with age, biological sex, and fatigue as fixed effects and participant as a random effect. To assess whether cellular changes impact bundle passive modulus, the average percent change in fiber passive modulus was correlated to average percent change in bundle passive modulus using a linear regression performed in SPSS. Of note, fibers were included from 2 older females not included in analyses for Figures 1-6 due to resulting inflation of the sample size for older females and subsequent effects of unequal group sizes on statistical analyses. However, inclusion of this subset in those analyses did not change the group means (data not shown).

## Results

### Participant anthropometrics and activity

11 young (6 females) and 11 older (7 females) adults were included in the analyses for this study. Older participants were aged 73 ± 4 years and younger participants were aged 21 ± 2 years. In the present sample of 22 participants, BMI was not significantly different between males and females (p=0.765) or between older and younger adults (p=0.219, Table 1). Height was significantly higher in males versus females (186.2 ± 11.5 vs. 162.9 ± 6.2, respectively, p<0.001) and was not different between older and younger participants (p=0.525). Weight was significantly higher in males versus females (81.2 ±10.8 vs. 65.4 ± 13.1, respectively, p=0.001) but was not different between older and younger adults (p=0.150). Although activity data were collected for most participants, the present analysis is missing activity data for 2 of the 4 older males due to equipment failure. The average number of steps completed per day was significantly higher in older (10402 ± 4463 steps/day) versus younger (7566 ± 1809 steps/day, p=0.008) participants. Time spent in moderate activity (75-125 steps/minute) was significantly higher in older (80.9 ± 37.7 minutes/day) versus younger (57.1 ± 16.1 minutes/day, p=0.014) adults. There were no significant differences in minutes spent in light (< 75 steps/minute, p=0.199) or vigorous (>125 steps/minute, p=0.084) activity between older and younger adults. There was no main effect of biological sex on step count (p=0.077), time spent in light activity (p=0.356), time spent in moderate activity (p=0.166), or time spent in vigorous activity (p=0.203).

**Table 1.** Anthropometric and activity data of older and younger participants.

| Age |  | N | BMI<br>(kg/m<sup>2</sup>) | Height<br>(cm)<sup>#</sup> | Weight (kg)<br><sup>#</sup> | Step Count<br>(steps/day) <sup>A</sup> | Light<br>Activity<br>(min./day) | Moderate<br>Activity<br>(min./day) <sup>A</sup> | Vigorous<br>Activity<br>(min./day) |
| --- | --- | --- | --- | --- | --- | --- | --- | --- | --- |
| Young | Male | 5 | 24.8 ± 1.2 | 184.2 ± 11.5 | 84.2 ± 8.8 | 7441 ± 1516 | 39.9 ± 7.8 | 54.0 ± 15 | 0.8 ± 1.1 |
|  | Female | 6 | 21.3 ± 1.0 | 160.9 ± 6.2 | 55.0 ± 2.8 | 7670 ± 2164 | 26.4 ± 8.1 | 59.7 ± 18.1 | 0.6 ± 0.6 |
| Older | Male | 4 | 23.0 ± 8.0 | 188.6 ± 27.2 | 77.5 ± 13.3 | 14936 ± 8625 | 36.3 ± 3.9 | 114.9 ± 70.7 | 5.0 ± 5.7 |
|  | Female | 7 | 27.6 ± 4.1 | 164.1 ± 6.5 | 74.4 ± 11.7 | 9106 ± 2312 | 41.6 ± 10.0 | 71.2 ± 23.8 | 1.6 ± 3.9 |

Symbols indicate significant difference between biological sex<sup>#</sup> or age<sup>A</sup> groups (p<0.05). Data are shown as mean ± standard deviation (SD).

### Fatiguing exercise and whole-muscle contractile performance

Average time to fatigue was not significantly different between older and younger adults (p=0.181) or males and females (p=0.067, Table 2). Similarly, fatigue ratio was not significantly different across age (p=0.817) or biological sex groups (p=0.201). Relative peak power was significantly higher in males versus females (6.08 ± 2.54 vs. 3.86 ± 1.49 W/kg, respectively, p=0.007) and higher in younger versus older adults (6.51 ±2.09 vs. 3.27 ± 0.84 W/kg, respectively, p<0.001), yet the interaction effect of age by biological sex on absolute peak power was not significant (p=0.774). Relative peak torque was significantly higher in males versus females (3.02 ± 0.91 vs. 2.07 ± 0.51 N/kg, respectively, p<0.001) and younger versus older adults (3.02 ± 0.76 vs. 1.90 ± 0.41 N/kg, respectively, p<0.001). The interaction effect of age by biological sex on relative peak torque was not significant (p=0.115).

**Table 2.** Fatigue data and whole-muscle performance of older versus younger adults.

| Age |  | Time to Fatigue (sec) | Fatigue Ratio | Relative Peak Power (W/kg) <sup>#A</sup> | Relative Peak Torque (N/kg) <sup>#A</sup> |
| --- | --- | --- | --- | --- | --- |
| Young | Female | 75±12 | 0.33±0.08 | 5.52±0.34 | 2.49±0.16 |
|  | Male | 88±39 | 0.37±0.16 | 7.49±2.69 | 3.64±0.73 |
| Older | Female | 75±10 | 0.32±0.06 | 2.67±0.21 | 1.70±0.39 |
|  | Male | 153±111 | 0.40±0.11 | 4.31±0.07 | 2.24±0.15 |

Symbols indicate significant difference between biological sex<sup>#</sup> or age<sup>A</sup> groups ( $p<0.05$ ). Data are shown as mean ± SD.

### Single fiber morphology and active mechanics

Fiber CSA was significantly different between males (0.0067 ± 0.0020 mm^2^) and females (0.0044 ± 0.0015 mm^2^, p<0.01) but was not different between older (0.0054 ± 0.0022 mm^2^) and younger (0.0055 ± 0.0019 mm^2^, p=0.857) adults (Table 3). Given the diversity in fiber-type distributions among the participant groups, statistical analyses were limited to MHC IIA and IIA/X fibers (n=257). Passive modulus was not different between MHC IIA and MHC IIA/X fibers at short or long lengths, in any participant group (data not shown). Active tension in this sample was not significantly different between non-fatigued and fatigued samples (p=0.080), males and females (p=0.410) or older and younger adults (p=0.569, Figure 1).

**Table 3.** Descriptive statistics of single fibers included in the study of effects of aging and fatigue on cellular passive mechanics.

|  |  | n | CSA (mm<sup>2</sup>) <sup>#</sup> | Length (mm) | Fiber type distribution (n) |  |  |  |
| --- | --- | --- | --- | --- | --- | --- | --- | --- |
|  |  |  |  |  | I | IIA | IIX | IIA/X |
| Young | Female | 107 | 0.0046 ± 0.0014 | 1.5 ± 0.4 | 23 | 50 | 2 | 24 |
|  | Male | 92 | 0.0066 ± 0.0019 | 1.7 ± 0.4 | 3 | 44 | 21 | 22 |
| Older | Female | 154 | 0.0043 ± 0.0016 | 1.5 ± 0.4 | 55 | 51 | 7 | 26 |
|  | Male | 92 | 0.0068 ± 0.0021 | 1.7 ± 0.5 | 60 | 29 | 2 | 11 |

Data are shown as mean ± SD. Percents are not reported for MHC I/IIA or I/IIA/IIX because they collectively made up only 6.2% of the overall dataset. Symbols indicate main effects of <sup>#</sup>biological sex ( $p<0.05$ ).

### Single fiber passive mechanics

Passive stress was analyzed at each SL studied throughout the passive stretch protocol. Passive Young’s Modulus was measured as the slope of the stress-strain curve at “short lengths” and “long lengths”. There were no main effects of age or biological sex on passive stress or Young’s Modulus when all fibers (non-fatigued and fatigued fibers) were considered.

The interaction of age and fatigue significantly affected passive stress at one SL (p=0.042 at SL 3.0 μm) and passive Young’s Modulus at short lengths (p=0.023). In the subset of non-fatigued fibers, age significantly affected passive stress at one SL (p=0.045 at SL 3.0 µm, Figure 2A) and passive modulus was higher in fibers from older adults versus younger adults at short lengths (p=0.042) but not long lengths (p=0.163, Figure 2B).

**Figure 2.**
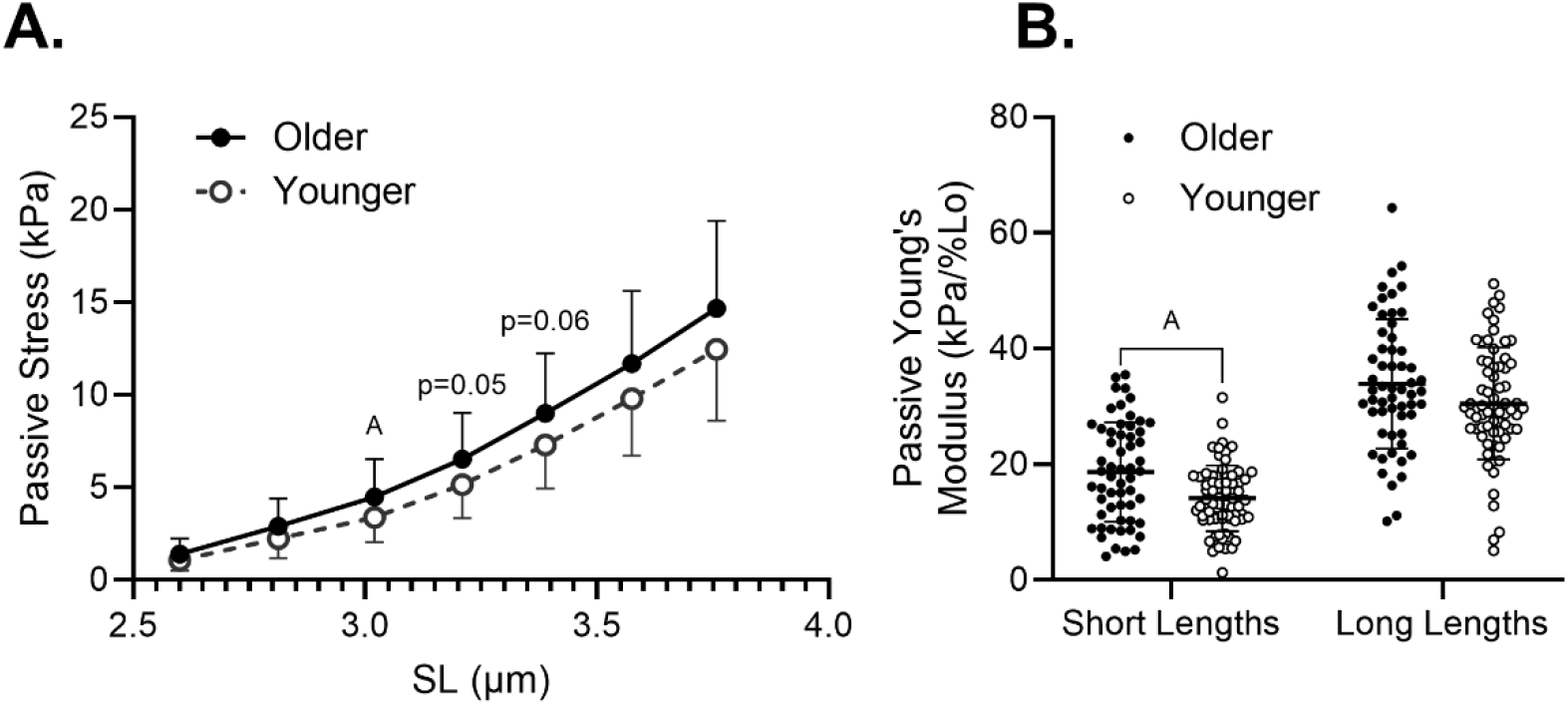
When only non-fatigued MHC IIA and MHC IIA/X fibers were considered, passive stress was significantly higher in fibers from older versus younger adults at SL=3.0 µm **(A)** and cellular passive modulus was significantly higher in fibers from older versus younger participants at short, but not long, lengths **(B)**. Data are shown as mean ± SD. ^A^ Indicates a significant effect of age (p<0.05).

When all fibers were considered, fatiguing exercise significantly reduced passive stress at all lengths (p<0.05 at SL 2.6-3.8 μm) and passive modulus at both short lengths (p<0.001) and long lengths (p=0.012). Furthermore, the interaction of fatigue and biological sex significantly affected passive stress at all SLs (p<0.01 at SL 2.6-3.8 μm) and passive Young’s Modulus at both short (p<0.001) and long (p=0.003) lengths, prompting further study of the effect of fatigue on passive mechanics in fibers from males and females, separately. As a result, it became clear that fatigue significantly reduces passive stress in fibers from males (p<0.01 at SL 2.6-3.8 μm, Figure 3A) but not females (Figure 3B). Similarly, fatigue-induced reductions in passive modulus were driven by males at short (p<0.001) and long (p<0.001) lengths, whereas modulus in single fibers from females was not significantly different with fatigue at short lengths (p=0.754) or long lengths (p=0.764, Figure 4).

**Figure 3.**
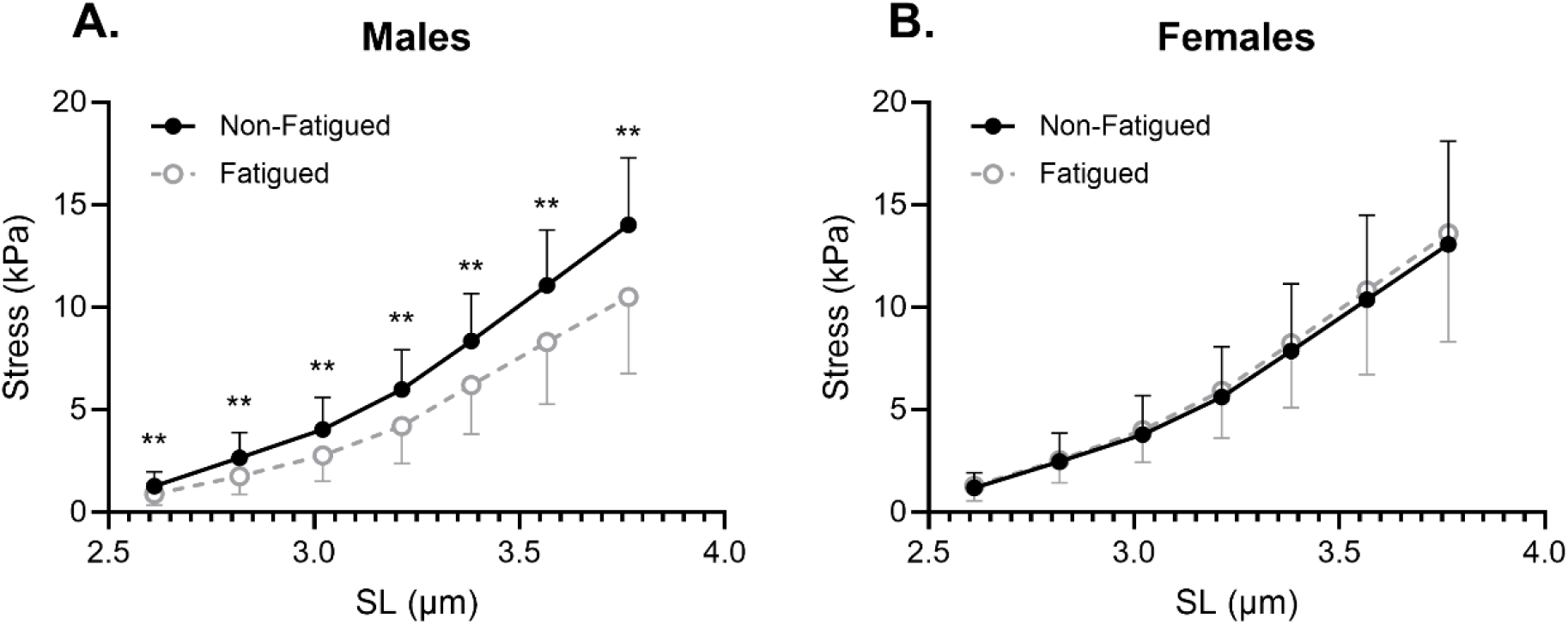
Cellular passive stress was significantly lower in fatigued versus non-fatigued fibers of males **(A)** but not in fibers of females **(B)**. Data are shown as mean ± SD. Symbols indicate a significant main effect of fatigue *(p<0.05) or **(p<0.01).

**Figure 4.**
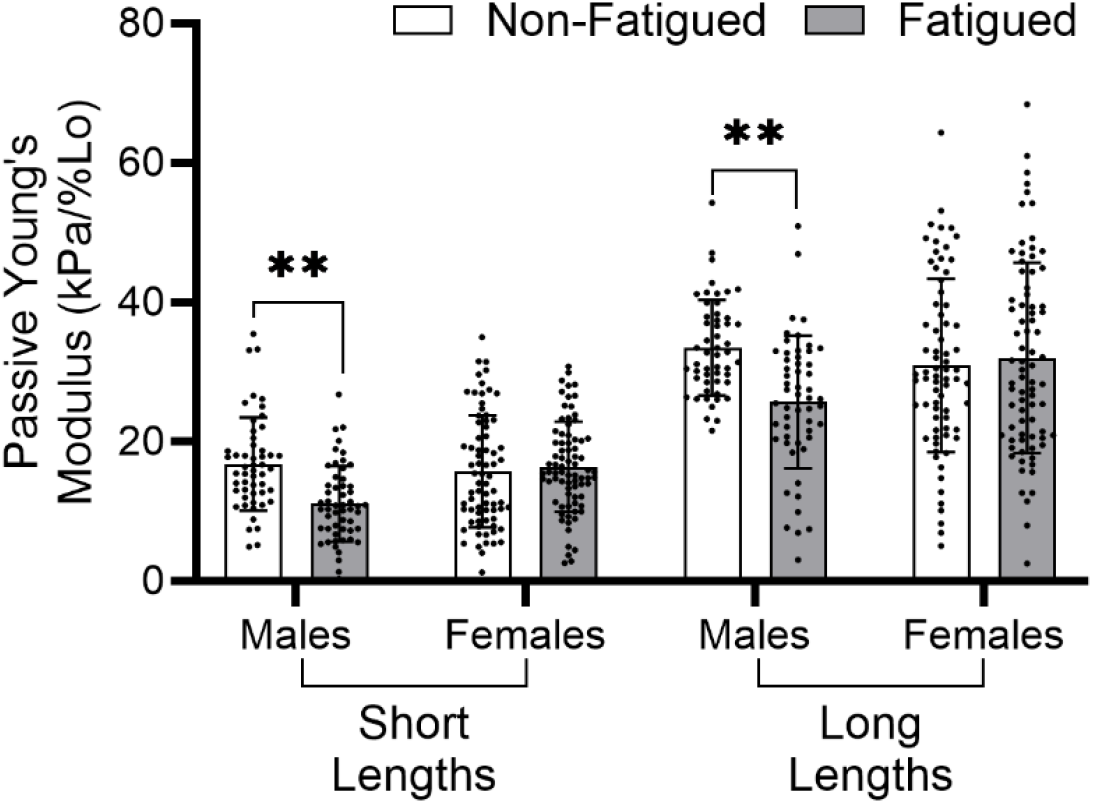
Cellular passive modulus is significantly reduced by fatigue at short and long lengths in fibers from males but not females. Data are shown as mean ± SD. Symbols indicate significant effects of fatigue, *(p<0.05) or **(p<0.001).

While there was no main effect of fatiguing exercise on cellular passive mechanics in female participants, inter-individual differences in cellular response were noted within the group. This observation prompted a qualitative look at changes to mean cellular passive modulus at short (Figure 5A) and long (Figure 5B) lengths in fatigued versus non-fatigued samples, for each participant. Whereas young and older males consistently demonstrate reductions in mean passive modulus, young and older female participants exhibit substantial variability in the effect of fatiguing exercise on mean passive modulus.

**Figure 5.**
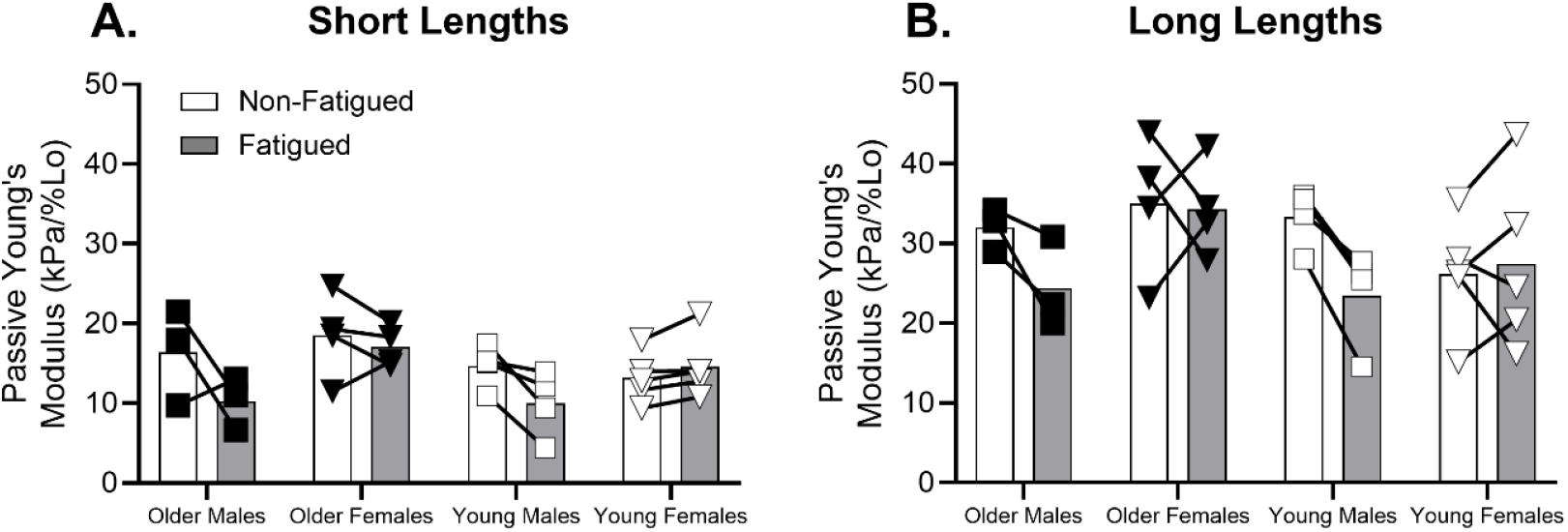
A qualitative look at the variability in change of mean cellular passive modulus on a per-individual basis. At short **(A)** and long **(B)** lengths, it appears that while male participants demonstrate consistent changes to passive modulus resulting from fatigue, females exhibit diversity in the response of passive modulus to fatiguing exercise. Filled square indicates older males, filled triangle indicates older females, open square indicates young males, open triangle indicates young females.

### Bundle sample characteristics

Passive stress and Young’s Modulus were measured in bundles from a subset of the participants included in cellular assays: 2 older males, 5 older females, 2 younger males, and 2 younger females. To account for the potential effect of variable fiber count and ECM quantity within each bundle on subsequent mechanical measures (Malakoutian *et al*., 2021), fiber to bundle ratios greater than 2 SD from the mean were excluded from analysis (n=8). Additionally, bundles producing passive modulus values greater than 2 SD from the mean (n=6) were considered outliers and excluded from analyses. Ultimately, 173 bundles were included in the present analysis (Table 4). The fiber to bundle ratio was not different by age (p=0.466), biological sex (p=0.845), or fatigue (p=0.800). Similarly, bundle length was not different by age (p=0.134), sex (0.143) or fatigue (p=0.699).

**Table 4.** Descriptive statistics of the bundles included these studies.

|  |  | N | Fiber to bundle ratio | Length (mm)<sup>#</sup> |
| --- | --- | --- | --- | --- |
| **Male** | Older | 19 | 13.3 ± 8.3 | 2.1 ± 0.4 |
|  | Younger | 18 | 10.0 ± 7.5 | 1.9 ± 0.3 |
| **Female** | Older | 80 | 19.2 ± 6.9 | 2.4 ± 0.5 |
|  | Younger | 56 | 12.1 ± 3.8 | 2.4 ± 0.5 |

Data are shown as mean ± SD. <sup>#</sup>Indicates a significant effect of biological sex (p<0.05).

### Bundle Passive Young’s Modulus

When all bundles were considered collectively, there were no significant main effects of fatigue (p=0.115), biological sex (p=0.769), or age (p=0.648) on bundle passive modulus. Given the relatively low numbers of participants within each group, the effect of fatigue on bundle passive modulus was explored in each participant group separately (Figure 6A). No significant effect of fatigue was found on bundle passive modulus in older males (p=0.184), older females (p=0.739), or younger males (p=0.831). However, bundle passive modulus was significantly higher in the fatigued versus non-fatigued bundles of young females (27.67±11.28 vs. 22.51±9.73 kPa/%Lo, respectively, p=0.033). A qualitative look at differences in mean bundle passive modulus in fatigued compared to non-fatigued samples for each participant (Figure 6B) highlights the variability in response of bundle passive modulus to fatiguing exercise across participant groups. To assess whether observations at the cellular level translate to the bundle level, the average percent change in fiber passive modulus was correlated to average percent change in bundle passive modulus (Figure 7). The resulting trend (p=0.10) for a positive correlation suggests that the effect of fatiguing exercise on cellular mechanics translates to higher levels of muscle organization.

**Figure 6.**
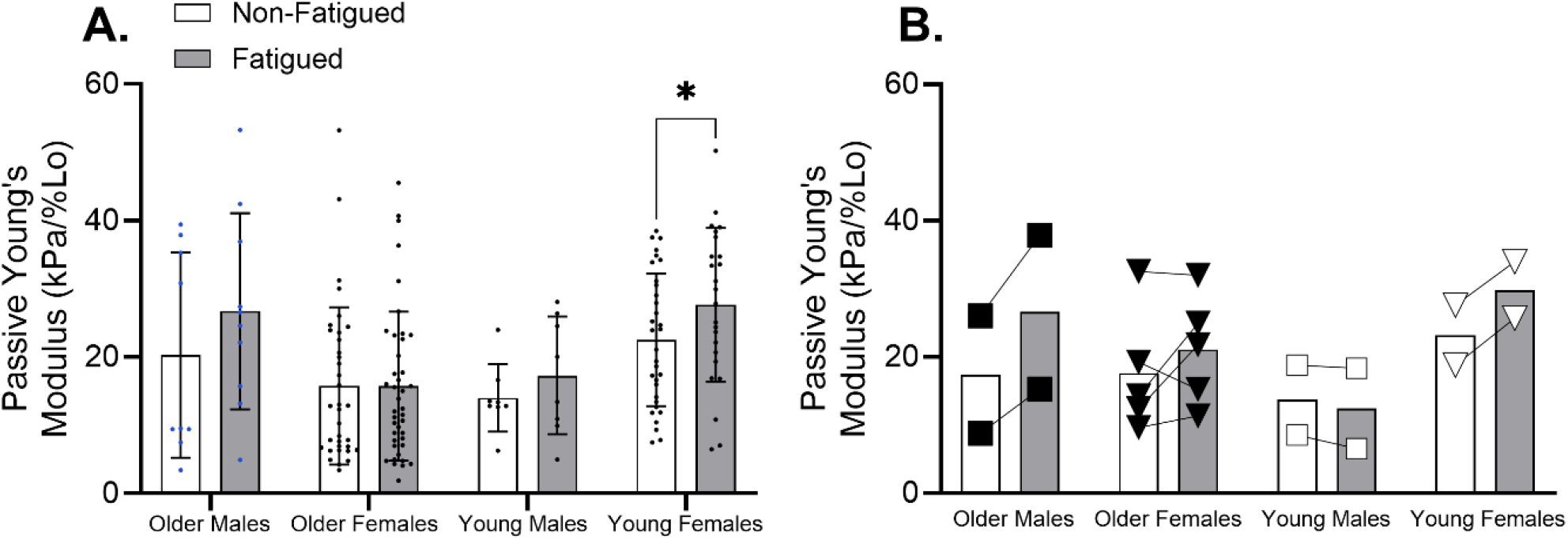
The effect of fatiguing exercise on bundle passive modulus was examined within each age and biological sex group. **(A)** Bundle passive modulus was significantly enhanced by fatiguing exercise in young females, but not in any other participant group. Data are shown as mean ± SD. * Indicates a significant effect of fatigue (p<0.05). **(B)** Visualization of relative fatigue-induced change in bundle passive modulus, presented on a per-individual basis, reveals considerable variability within and between groups. Filled square indicates older males, filled triangle indicates older females, open square indicates young males, open triangle indicates young females.

**Figure 7.**
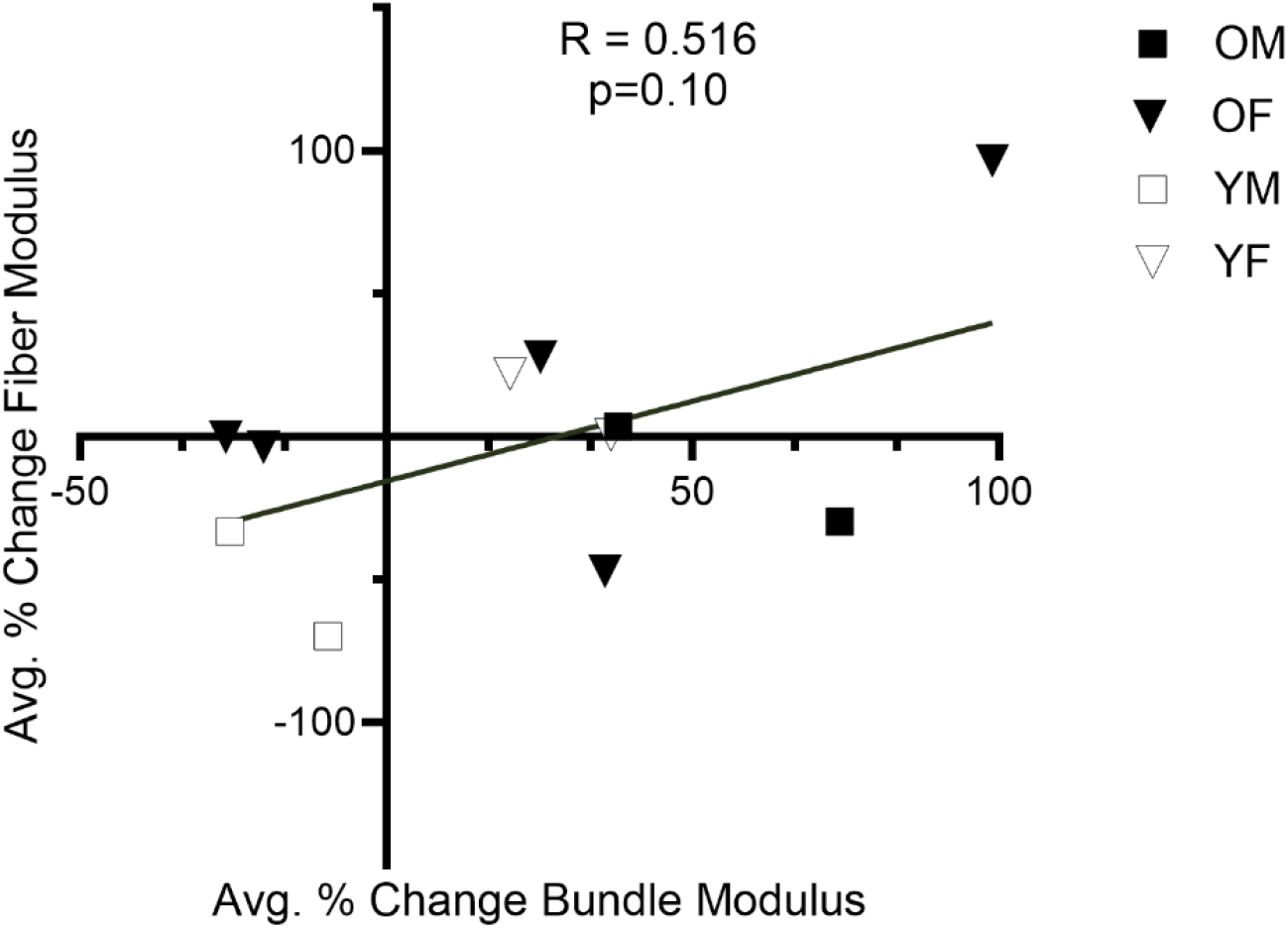
A modest trend (p=0.10) for association between fatigue effects at the fiber and bundle level suggests that changes to cellular passive modulus may influence passive modulus at higher levels of skeletal muscle organization. OM = older males, OF = older females, YM = young males, YF = young females.

## Discussion

In the present study, we report male-specific reductions in passive Young’s Modulus with fatigue and inconsistent responses in females, regardless of age. These results contradict our initial hypothesis that modulus in older adults would be resistant to the acute effects of fatiguing exercise, regardless of sex. These findings extend our previous work characterizing the effect of fatiguing exercise on single fiber passive mechanics in young adults (12, 13) to include healthy older adults and show durability of this response in males with variability in responses across female populations regardless of age.

### Study Sample

In this cohort, older participants had greater step count per day, and more time spent in moderate physical activity per day, than younger adults (Table 1). Given the tendency for physical activity to remediate other age-related changes in muscle (25), the high activity level of older participants in this study may have obscured age-related effects on passive muscle mechanics that may exist otherwise. Notably, whole-muscle performance from the present sample of older and younger adults was reduced in the aged group (Table 2), consistent with previous reports (26). However anthropometric measures were not different across age groups and at the cellular level there was no difference in fiber CSA (Table 3), agreeing with some (16, 18) but not all (27) previous reports. This lack of a fiber size difference suggests the comparatively high physical activity level of the older participants protected them from some age-related changes to muscle morphology and, potentially, function. We previously observed that chronic resistance training not only alters cellular passive stress and modulus, but also mediates the effect of fatiguing exercise on these cellular passive measures (13). However, MHC II fibers appear less well protected from the effects of aging by physical activity (28). Nevertheless, the influence of activity differences in this cohort must be considered when interpreting the present results. Despite the potential influence of differences in physical activity across age groups, single fiber active isometric tension (Figure 1) was not significantly different by biological sex or participant age, agreeing with previous reports (12, 29).

### Aging increases passive stress and Young’s Modulus at short sarcomere lengths

We hypothesized that muscle fibers from older adults would exhibit higher passive stress and modulus compared to fibers from younger adults. Our measures revealed age-related increases in passive stress (Figure 2A) and Young’s Modulus at short sarcomere lengths (Figure 2B), partially supporting our hypothesis. This result suggests age-related increases in cellular passive mechanics are limited to short sarcomere lengths and may not extend to long sarcomere lengths, where thin/thick filament overlap is limited and titin is the predominant contributor to cellular passive modulus. This observation was surprising, considering that fibers from older adults generated increased passive stress in SL > 3.0 µm in prior studies (16, 18). An important consideration when comparing our results to those in the existing literature is that our analyses were restricted to MHC IIA and IIA/X fibers while others were pooled, though MHC isoform has not been definitively shown to impact passive modulus. Additionally, physical activity has been demonstrated to preserve skeletal muscle function in aging (30–32). Therefore, the elevated physical activity levels of the older participants in this study may have attenuated expected age-related changes to skeletal muscle and limited our ability to test the effect of aging per se on cellular passive mechanics.

### Biological Sex, but not Aging, Mediates the Effect of Fatiguing Exercise on Cellular Passive Stress and Young’s Modulus

We hypothesized that aging would mitigate the effect of fatiguing exercise on cellular passive stress and Young’s Modulus. This hypothesis was based on the notion that older adults experience reduced intramyocellular perturbation (e.g. pH, inorganic phosphate) during fatiguing exercise than their younger peers (33–35), therefore the accumulation of potential modulators of cellular stiffness via post-translational modifications to titin would be less in muscle from older versus younger adults. Surprisingly, the present results suggest comparable magnitudes of change in older and younger adults (Figure 5). In the present study, fatigue was quantified as fatigue ratio, the ratio of final power to initial power produced during the fatiguing leg extension exercise (Table 2). A similar fatigue ratio across the age groups is at least in part reflective of comparable intramyocellular perturbations due to fatiguing exercise, which may contribute to the lack of an age effect on magnitude of change to cellular passive mechanics following fatiguing exercise.

A strong interaction between biological sex and fatigue was found to impact cellular passive mechanics, consistent with our previous results reported in young adults (12, 13). Whereas younger and older males demonstrated clear reductions in passive stress (Figure 3A) and Young’s Modulus (Figure 4) in fatigued versus non-fatigued fibers, these measures were not significantly different following fatiguing exercise in females (Figure 3B, Figure 4). Furthermore, the lack of a significant effect of fatigue on cellular passive mechanics in females appears to derive from high variability in response to the fatiguing exercise across participants at short (Figure 5A) and long lengths (Figure 5B). We previously speculated that this biological sex effect may reflect inter-individual differences in circulating sex hormones such as estrogen amongst female participants (12). However, this idea was not supported by the comparison of passive stress and Young’s Modulus in this cohort of older versus younger participants. In females, estrogen levels decline in the years before menopause, after which they remain constantly low (36). In males, testosterone levels begin dropping around 40 years of age and continue to decline at a constant rate thereafter (37). Therefore, it would be expected that any effect of sex hormones on titin mechanics would be evident in the comparison of cellular passive mechanics of older versus younger participants, given the anticipated differences in hormone profiles. However, in the present sample there was no effect of aging on the acute regulation of cellular passive mechanics, suggesting that alternative mechanisms must be considered.

We have previously proposed heat shock protein (HSP) 27 and alphaβ-crystallin as alternative mechanisms of altered cellular passive mechanics following fatiguing exercise in males versus females. Interestingly, the quantity of these small HSPs has been demonstrated to increase with advanced aging in human vastus lateralis muscle (38), the same muscle that was biopsied for the present study. In line with the hypothesis that HSP 27 and alphaβ-crystallin interact with titin (39) to reduce titin-based stiffness in a sex-dependent manner (40), the age-related increases in small HSP concentration may preserve the acute regulation of cellular passive mechanics in older males, despite concurrent age-related changes to sex hormones (37). Measurement of small HSP was not performed in the present study but should be considered in future investigations of chronic and acute mediators of cellular passive mechanics.

### Extending cellular measures to the tissue level

We hypothesized that non-fatigued fiber bundles would exhibit higher passive stress and modulus in samples from older versus younger participants, based on reports of increased ECM collagen content and cross-linking in advanced age (17, 41). In the present results, there was no main effect of age on bundle passive modulus (Figure 6A). We also hypothesized that fatiguing exercise would reduce passive stress and modulus at the bundle level in older and younger adults. However, the present results do not support this hypothesis. While passive modulus was significantly higher in bundles from the fatigued versus the non-fatigued limb in samples from young females, a significant effect of fatigue on passive modulus was not observed in any other participant group. The high variability across samples within each participant group (Figure 6B) may have impaired our ability to detect an effect of fatiguing exercise on bundle mechanics. Alternatively, the high activity level present in this cohort of older adults may have mediated any age-related changes in skeletal muscle structural (ECM) or proteomic (titin) changes expected of aged skeletal muscle.

Importantly, bundle size impacts the passive mechanical measures of the sample, presenting an experimental limitation to this study. Specifically, larger bundles tend to have lower passive modulus, likely due to the inclusion of relatively less ECM (42). ECM is stiffer compared to single fibers, which explains the non-linear increase of passive mechanical properties from single fiber to fascicle and whole-muscle scales of organization (43). Previous work (44) suggested that smaller fibers are stiffer compared to larger fibers due to the greater proportion of the stiffer basement membrane compared to the relatively less stiff internal area of the fiber. Coupled with the notion that smaller fibers within a bundle increase the relative proportion of CSA that is occupied by ECM (42), it is possible that differences in fiber size and bundle size may have contributed to unequal inclusion of ECM across samples and subsequent variability in bundle passive stress and modulus measures. However, an estimate of the number of fibers included per bundle (“fiber to bundle ratio”) suggests some consistency across sample groups (Table 4).

Despite the high degree of variability in bundle-level passive modulus data, a correlation analyses between fatigue-induced changes in modulus at the fiber level and corresponding changes in fiber bundles on a per-individual basis (Figure 7) suggest a positive association between fatigue induced changes in modulus at these scales of muscle tissue composition. This correlation may indicate that changes to single fiber mechanics influence higher levels of skeletal muscle organization, but low sample size and a lack of statistical significance preclude drawing strong conclusions.

## Conclusions

We find significant fatigue-induced reductions in passive stress and Young’s modulus in males that do not manifest in females, suggesting that fatiguing exercise does not impact cellular passive mechanics in female participants. However, examination on a per-individual basis supports that females exhibit a more diverse response of cellular mechanics to fatiguing exercise. Further investigations into the source of this sex-based variability will be especially important given the greater propensity for frailty and falls risk in older adult females (45). Our attempt to translate fiber-level data to the tissue level demonstrated no significant effect of fatiguing exercise, though variance in bundle mechanics appeared to derive from the inherent characteristics of the constituent fibers, rather than from ECM-based mechanisms. Together, these results support the notion that biological sex mediates the effect of fatiguing exercise on skeletal muscle passive mechanics in younger and older adults. Whether cellular changes to passive mechanics manifest at higher levels of muscle organization remains unclear. This work contributes to efforts aimed at understanding how skeletal muscle mechanical properties are regulated, which may have implications for reducing falls-risk in older adults.

## Data availability statement

The raw data that support the findings of this study are available from the corresponding author upon reasonable request.

## Acknowledgements

The Authors wish to thank the volunteers for their invaluable contributions to our work. Additionally, thanks are due to Cameron Mulder of the Knight Library at the University of Oregon for valuable statistical consultation.

## Funding

This research was supported by the Wu Tsai Human Performance Alliance and NIH R21AG077125-01A1.

## Disclosures

The authors declare no competing interests, financial or otherwise.

## Author Contributions

These experiments were conducted in the Muscle Cellular Biology Laboratory in the University of Oregon Human Physiology Department. G.E.P., J.O.D., A.W.R., K.W.N, and D.M.C. designed and performed experiments. G.E.P., J.O.D., A.W.R., K.W.N, and D.M.C. analyzed and interpreted study data and revised the manuscript. G.E.P., J.O.D., and D.M.C. drafted the manuscript. D.M.C. conceived of and directed the study. All authors approved the final version of the manuscript and agree to be accountable for all aspects of the work in ensuring that questions related to the accuracy or integrity of any part of the work are appropriately investigated and resolved. All designated as authors qualify for authorship and all those who qualify for authorship are listed.

## Notes

### Competing Interest Statement

The authors have declared no competing interest.

